## Supplemental Figures 1-7 for "Embedding muscle fibers in hydrogel improves viability and preserves contractile function during prolonged *ex vivo* culture"

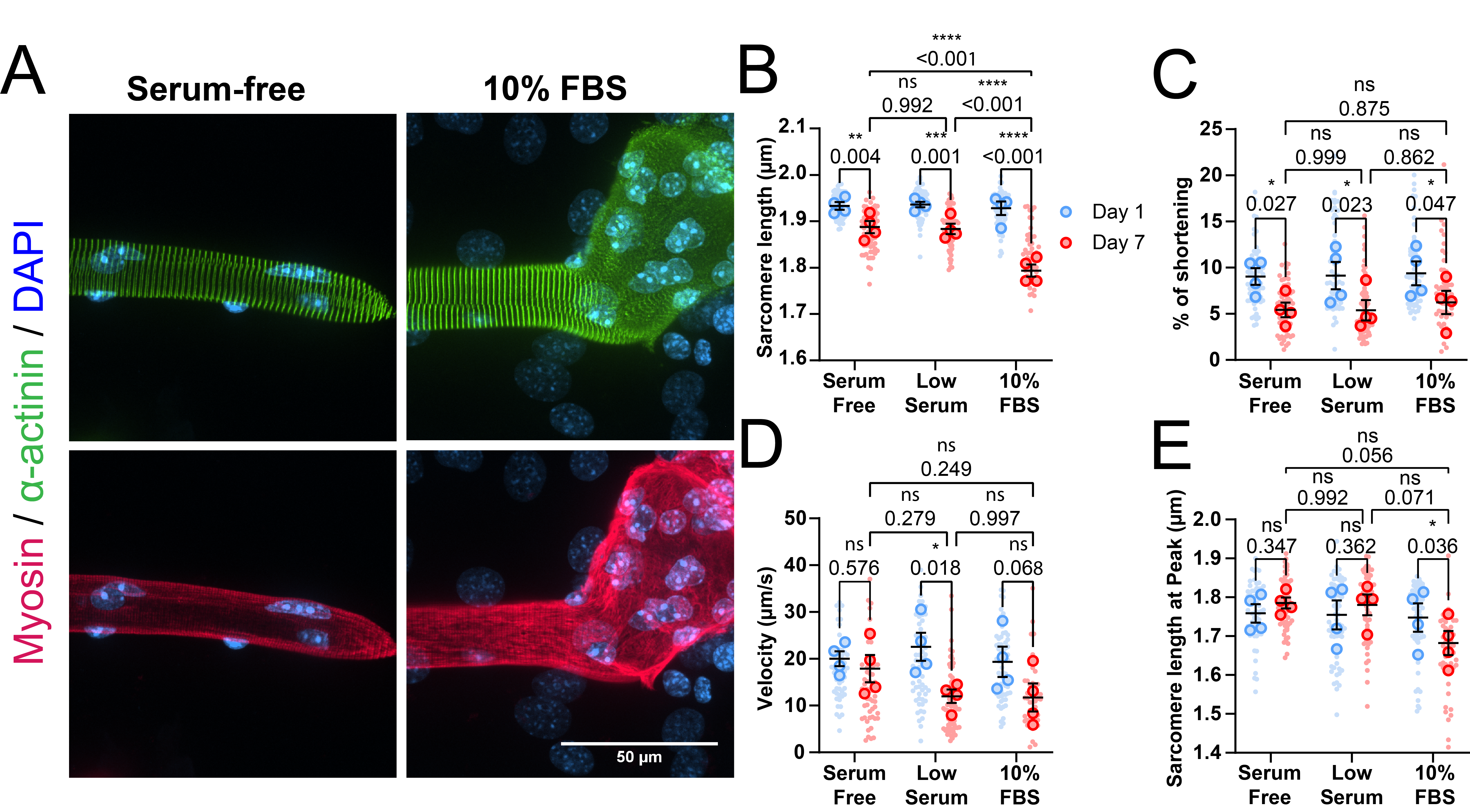

**Supplemental Figure 1:** **Effects of serum in culture medium on muscle fiber dedifferentiation.** (A): Representative confocal maximum intensity projection images immunostained for α-actinin (green), myosin (red) and nuclei (blue) in muscle fibers cultured for 7 days in Serum-free medium and medium containing 10% FBS. Note the lack of sarcomere misalignment in the muscle fiber cultured in Serum-free medium and the large dedifferentiated mass in the muscle fiber cultured in 10% FBS. (B-E): Contractile measurements of muscle fibers cultured in low serum, medium containing 10% FBS and serum-free medium, measured at Day 1 and Day 7 of culture. (B): Quantification of resting sarcomere length at Days 1 and 7 ex vivo culture. (C): Quantification of percentage of sarcomere shortening at Days 1 and 7 ex vivo culture. (D): Quantification of contractile velocity at Days 1 and 7 ex vivo culture. (E): Sarcomere length during maximal contraction of muscle fibers kept in culture. Data are means ± SEM N = 4 for mice and n = fibers. Significance was determined using a two-way ANOVA with p < 0.05 considered as significant with * = p < 0.05, ** = p < 0.01, *** = p < 0.001.

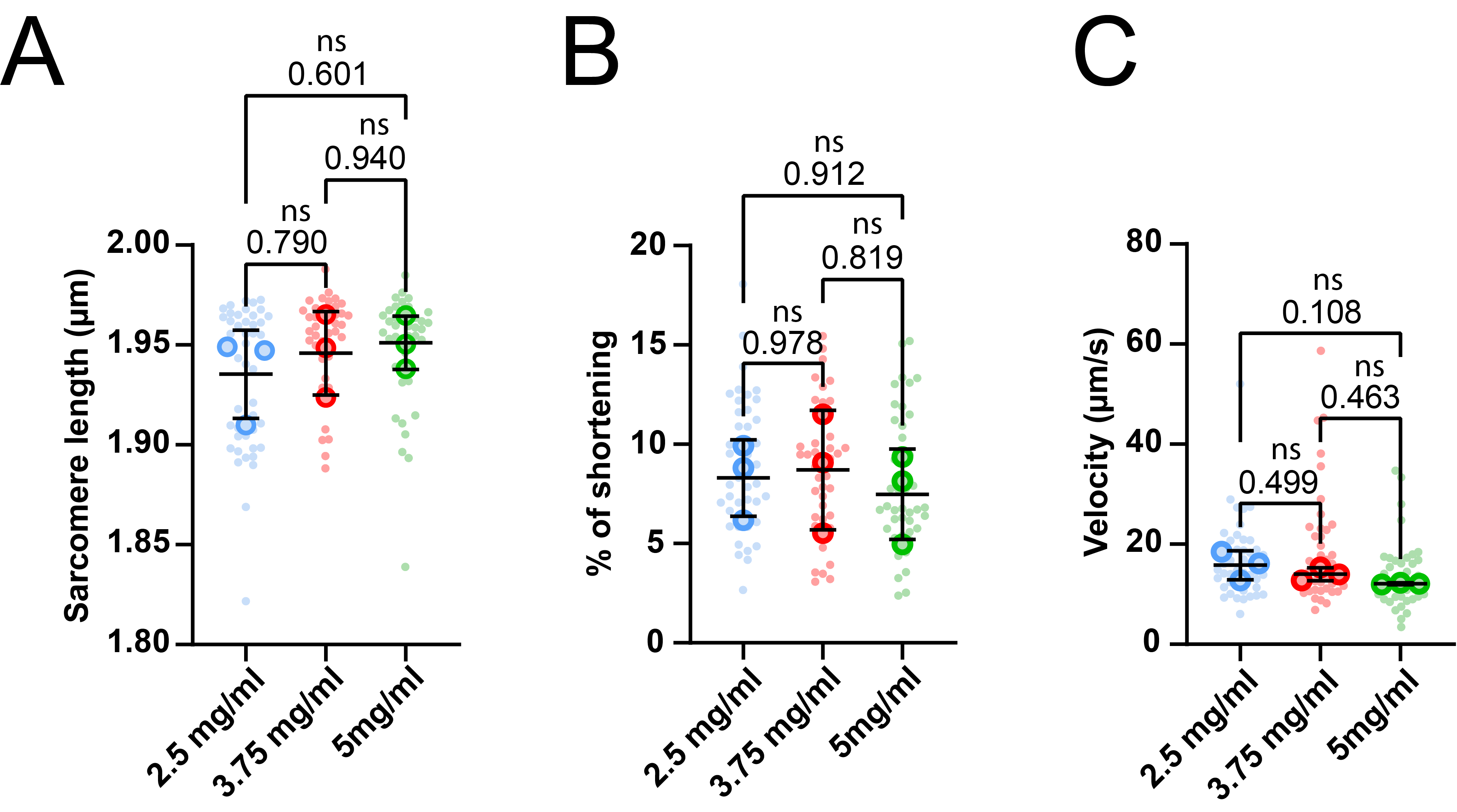

**Supplemental Figure 2: Contractile measurements of muscle fibers in increasing 3D hydrogel concentrations, measured at Day 1 of culture.** (A): Quantification of resting sarcomere length of muscle fibers in 3D hydrogel. (B): Quantification of sarcomere shortening of muscle fibers kept in 3D hydrogel. (C): Quantification of the contractile velocity of muscle fibers kept in 3D hydrogel. Data are means ± SEM N = 3 for mice and n = fibers. Significance was determined using a one-way ANOVA with p < 0.05 considered significant.

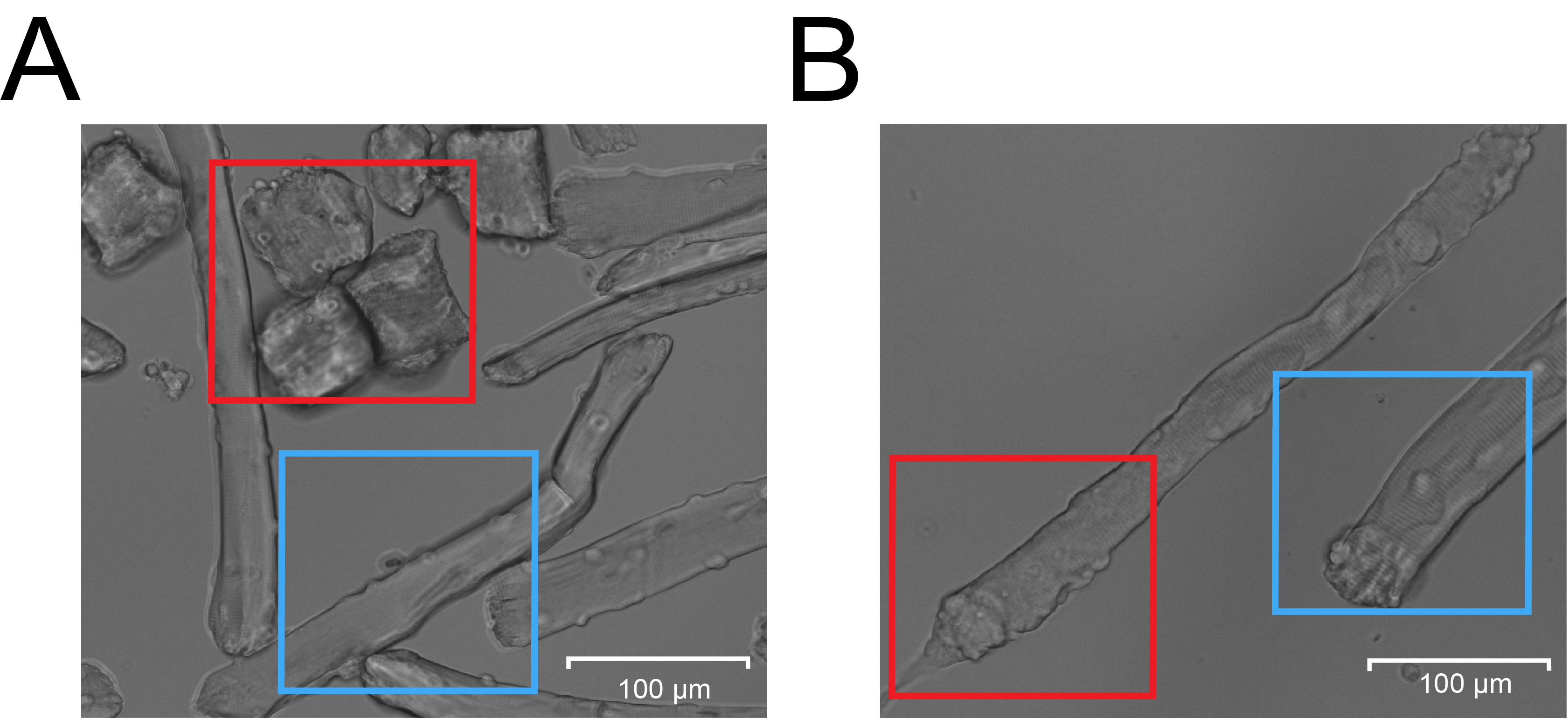

**Supplemental Figure 3: Muscle fiber classification examples** (A): Example image used for the classification of muscle fiber viability in which the red box includes dead/hypercontracted muscle fibers and the blue box includes living muscle fibers. (B): Example image used for the classification of muscle fiber dedifferentiation, in which the red box indicates a dedifferentiated muscle fiber and the blue box indicates a normal muscle fiber.

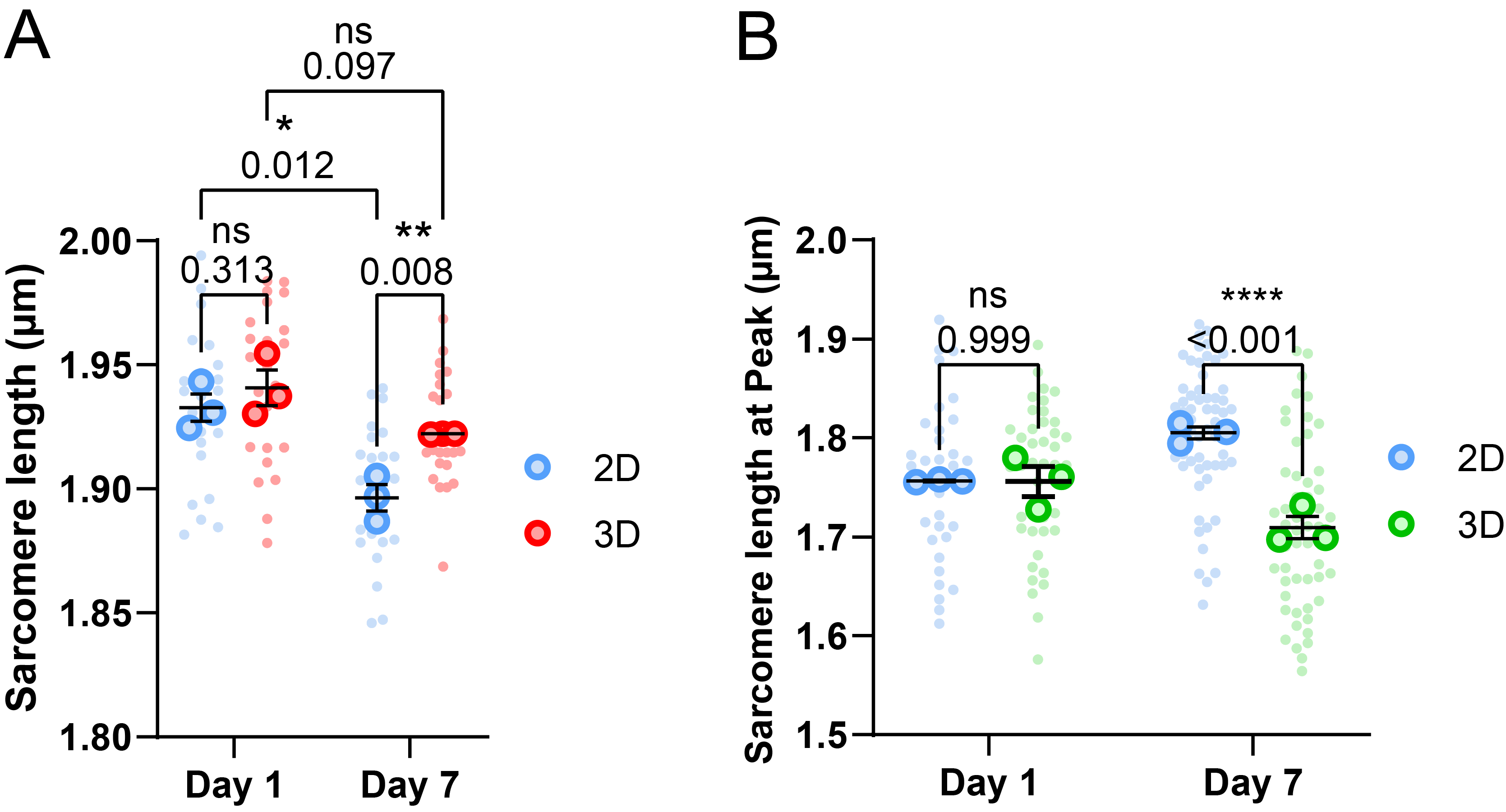

**Supplemental Figure 4: Sarcomere length measurements of muscle fibers cultured in 2D and 3D culture systems at Day 1 and Day 7 of culture.** (A): Quantification of resting sarcomere length of muscle fibers in 2D and 3D tracked over 7 days, which was used for normalization of muscle fiber width measurements in **Figs. 1D** and **2F**. (B): Sarcomere length at peak contraction of muscle fibers cultured in 2D and 3D, at Day 1 and Day 7. Data are means ± SEM N = 3 for mice and n = fiber. Statistics performed using a repeated measure two-way ANOVA in panel A and two-way ANOVA in panel B, with p < 0.05 considered as significant with * = p < 0.05, ** = p <0.01 and **** = p < 0.001.

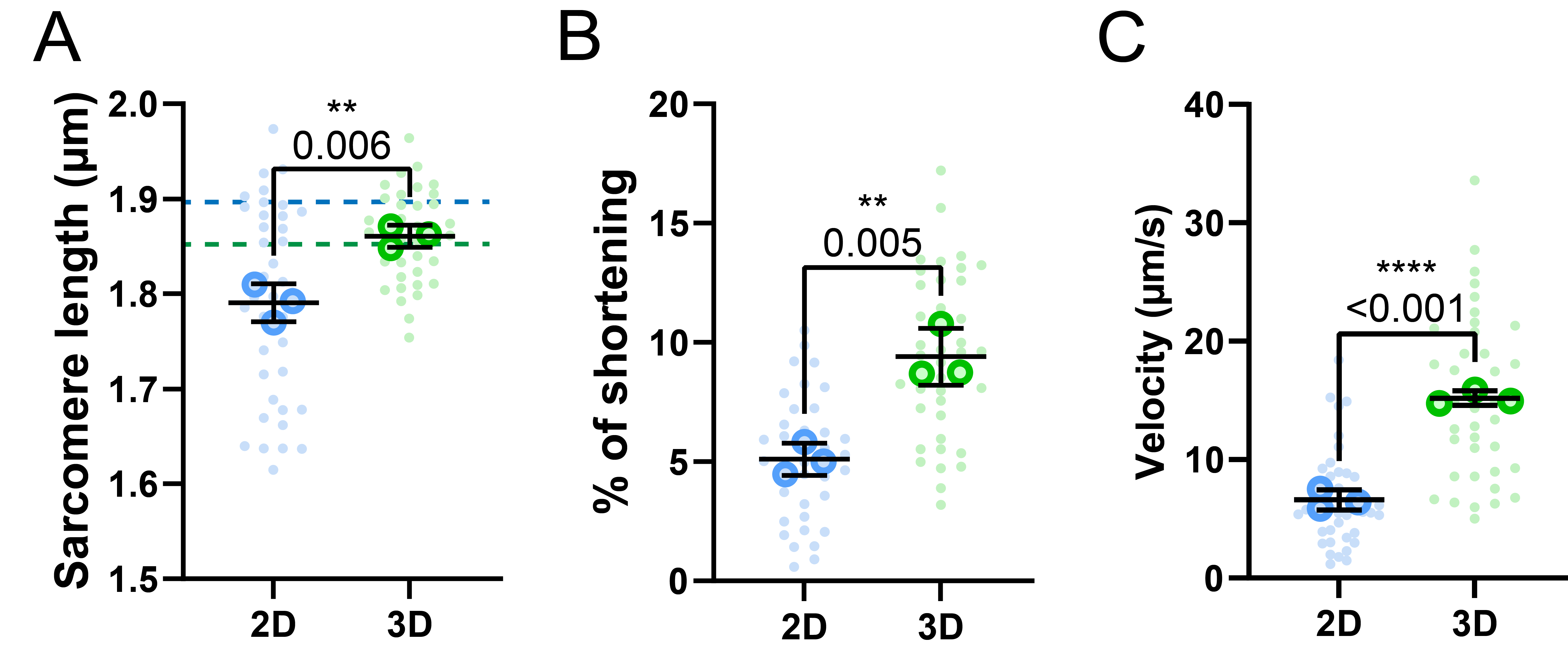

**Supplemental Figure 5: Contractile measurements of muscle fibers cultured in 2D and 3D, measured at Day 10 of culture.** (A): Quantification of resting sarcomere length of muscle fibers cultured for 10 days. Dashed line shows resting sarcomere length values of muscle fibers cultured for 7 days presented in Fig. 2. Note that resting sarcomere continues to decrease in 2D, while it is maintained in 3D (B): Quantification of sarcomere shortening of muscle fibers cultured for 10 days. (C): Quantification of the contractile velocity of muscle fibers cultured for 10 days. Data are means ± SEM N = 3 for mice and n = fibers. Significance was determined using student’s t-test with *p* < 0.05 considered as significant with ** = *p* < 0.01 and **** = *p* < 0.0001.

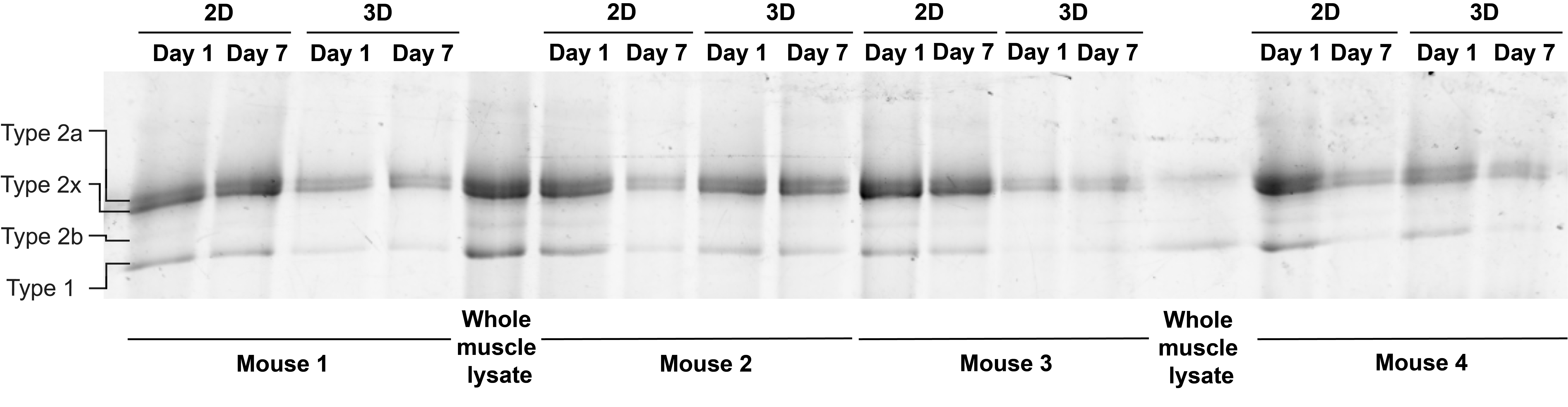

**Supplemental Figure 6: SDS-PAGE gel for identification of myosin heavy chain composition.** SDS-PAGE separation of myosin heavy chain isoforms from pooled FDB muscle fibers of four mice cultured in 2D and 3D, on Day 1 and Day 7. Mixed whole muscle lysate from mouse Soleus and EDL was used as a control in lanes 5 and 14.

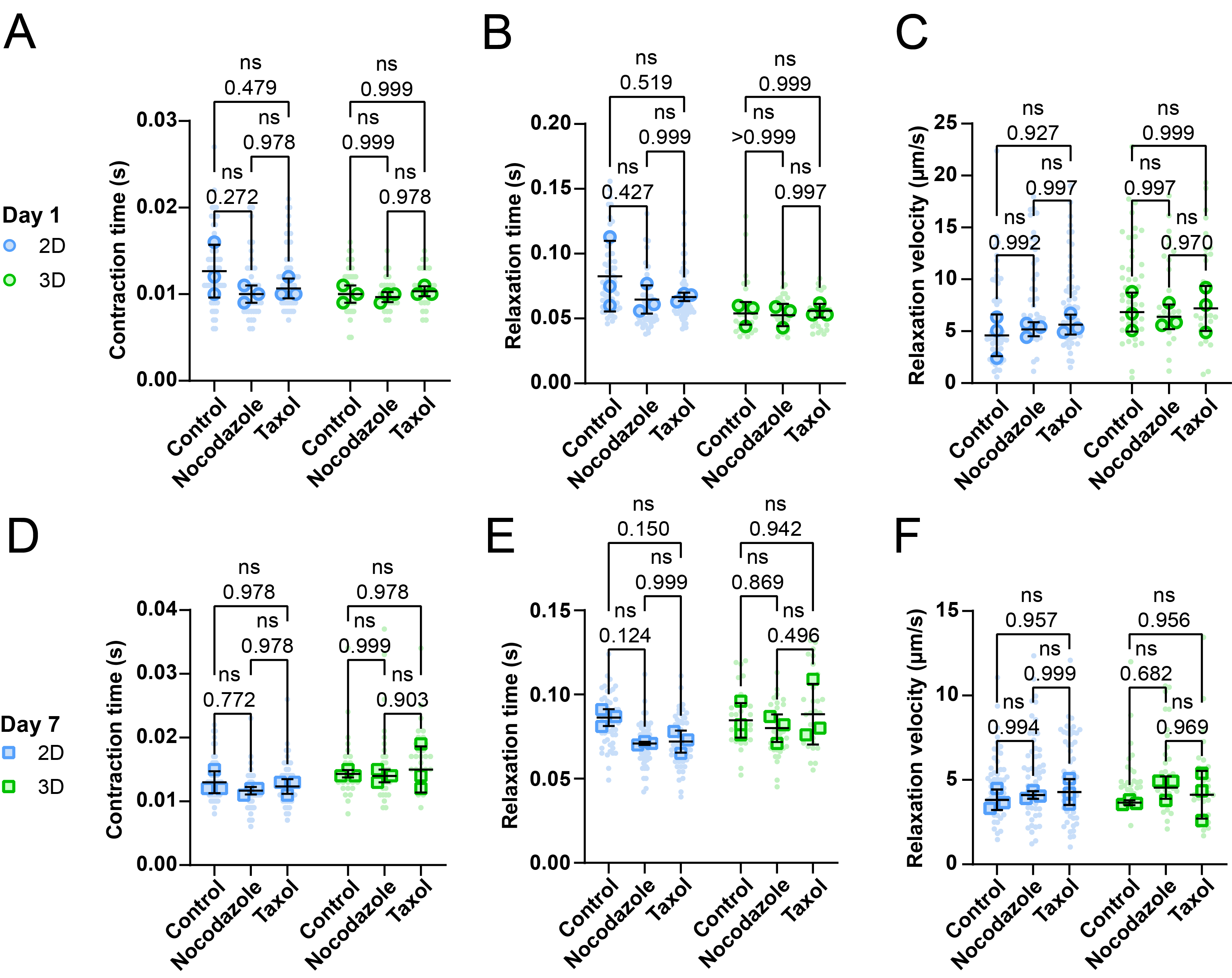

**Supplemental Figure 7: Contractile parameters in response to microtubule network manipulation in muscle fibers change during *ex vivo* long-term cultivation**(A-C): Quantification of contraction, relaxation duration and relaxation velocity measurements of muscle fibers cultured in 2D and 3D at Day 1, treated with Nocodazole, Taxol or untreated (control). (D-F): Quantification of contraction, relaxation duration and relaxation velocity measurements of muscle fibers cultured in 2D and 3D at Day 7, treated with Nocodazole, Taxol or untreated (control). Data are means ± SEM; *N* = 3 mice and *n* = fibers. Significance was determined using one-way ANOVA within culture condition (2D or 3D) with *p* < 0.05 considered as significant.
